# Synchrotron XRD based Structural analysis of Novel Fleroxacin Cocrystals synthesized using coformers

**DOI:** 10.1101/2025.07.16.664812

**Authors:** Ilham Zaky Wilson, J. Steven Swinnea, Leela Raghava Jaidev Chakka, Mohammed Maniruzzaman

**Affiliations:** PharmE3D Lab, School of Pharmacy, The University of Texas at Austin, Austin, TX 78756, USA; Cockrell School of Engineering, The University of Texas at Austin, Austin, TX 78756, USA; PharmE3D Lab, School of Pharmacy, The University of Mississippi, Oxford, MS 38655, USA

**Keywords:** Cocrystals, Synchrotron XRD, Fleroxacin, Powder X-ray Diffraction, Crystallization, Ball Milling, Antibiotic, Small Molecule, Material Science, Carboxylic Acid

## Abstract

Cocrystals have emerged as an exciting avenue in the pharmaceutical industry for altering the behavior of drugs while preserving the molecular structures of the active pharmaceutical ingredients (APIs). However, the preparation of pharmaceutical cocrystals remains relatively uncommon, presenting a potential opportunity for innovation. In this study, we developed cocrystals of Fleroxacin, a fluoroquinolone antibiotic belonging to the quinolone antibiotic class used in treating bacterial infections. Our approach involved co-crystallizing Fleroxacin with coformers (nicotinamide, salicylamide, and acetaminophen). The combination of Fleroxacin with different coformers allows us to explore chemical properties and the crystal efficacy. To achieve cocrystal formation, we employed a novel catalyst, pure glacial acetic acid, in conjunction with a ball milling machine. This methodology is particularly notable as it represents a first-time application of pure glacial acetic acid for co-crystallization and the co-crystallization of Fleroxacin. The cocrystals characteristics were analyzed using Synchrotron Powder X-ray Diffraction. The results showed the formation of novel cocrystals of Fleroxacin. The findings of this study contribute to the expanding body of knowledge on co-crystallization techniques and their potential applications in pharmaceutical development, especially for carboxylic acid-based drugs and drugs with very poor water solubilities but great permeabilities.

## 1. Introduction

Defined as crystalline solids made from at least two, and sometimes three^1^ molecules (or ions)^2^, ideally stoichiometric ratio^3^ cocrystals have a long-standing history dating back to the 19th century when Friedrich Wöhler studied the first reported cocrystal, quinhydrone in 1844^4^. Over the following decades and centuries, numerous cocrystals were discovered and reported in Paul Pfeiffer’s work “Organische Molekulverbindungen” published in 1922^4^. Cocrystals can be categorized as organic-organic or organic-inorganic compounds, contributing to their diverse chemical nature^4^. Although cocrystals represent only about 0.5% of the crystal structures archived in the Cambridge Structural Database (CSD)^5^, their importance cannot be overlooked. Cocrystals have proven to be valuable in various sectors, encompassing pharmaceuticals, textiles, paper, chemical processing, photography, propulsion, and electronics^4^.

One of the major advantages of cocrystals is that they can be easily synthesized using standard crystallization techniques such as evaporation and melting^6,7^. Even sublimation has been reported^6^. This eliminates the need for extensive research and development of new materials (hence, low-risk high-reward)^6^. In fact, significant advancements have been achieved, particularly in the field of explosives research^8,9^. Chiefly, the explosive cocrystal papers mainly discuss about transferring a beneficial property of a co-explosive to another explosive^10^. For example, Matzger *et. al*. has made three melt-castable energetics: 2,4,6-trinitrotoluene (TNT), 2,4-dinitroanisole (DNAN), and 1,3,3-trinitroazetidine (TNAZ) even more shock sensitive by cocrystallization with cyanuric triazid^10^. Therefore, it makes sense to wonder if this approach can be extended to pharmaceuticals: transferring desirable properties such as solubility to a less or even insoluble drug.

As mentioned before, considering the transformative effects of cocrystals in various fields (at least in explosive research), it is reasonable to speculate whether their properties can be manipulated to enhance the solubility of APIs. At the same time, interestingly in the pharmaceutical industry, most drugs candidates fail to commercialize due to bad biopharmaceutical properties instead of toxicity or lack of efficacy^11^. Solubility plays a crucial role in the bioavailability of drugs^12^ since the effectiveness of a drug relies heavily on its solubility. The chief reason is that most drugs are solids under room temperature and pressure conditions and delivered orally^13^. The oral route involves dissolution of the drug in the gastrointestinal (GI) tract then permeation into the bloodstream^13^. Particularly for those classified as Biopharmaceutics Classification System (BCS) Class II drugs, which exhibit high permeability but poor water solubility^14,15^. If a drug substance cannot dissolve, it cannot be absorbed, resulting in reduced bioavailability^12^. Therefore, it makes sense to consider enhancing solubility through co-crystallization with highly water-soluble coformers, preferably nutrients or well-tolerated over-the-counter painkiller drugs. In fact, there have been many attempts to increase the solubility of BCS Class II & IV drugs showing that this is indeed a valid concern^16,17^. Motivated by these prospects, our study focuses on the synthesis and characterization of cocrystals of Fleroxacin, a versatile fluoroquinolone antibiotic belonging to the quinolone antibiotic class.

Fleroxacin (see figure 1A(I)) has shown effectiveness in treating various bacterial infections and falls under the BCS Class II, category with negligible water solubility^15^. However, during our screening and experimentation for this project, we had discovered that this drug exhibits exceptionally high solubility in glacial acetic acid, forming solutions without any suspended particles. Despite the extensive research on cocrystals, as of 2023, no known cocrystal of Fleroxacin has been reported in the literature. The closest example is an abstract of a presentation by Changquan Calvin Sun from the University of Minnesota, who obtained single crystals of the Fleroxacin-saccharin cocrystal through slow evaporation of an acetonitrile solution (as of Apr 2024, no journals from that abstract are known to our knowledge)^18^. Our study distinguishes itself from previous work by employing ball milling as the synthesis method, aiming for gram-scale production and utilizing novel catalysts.

**Figure 1:**
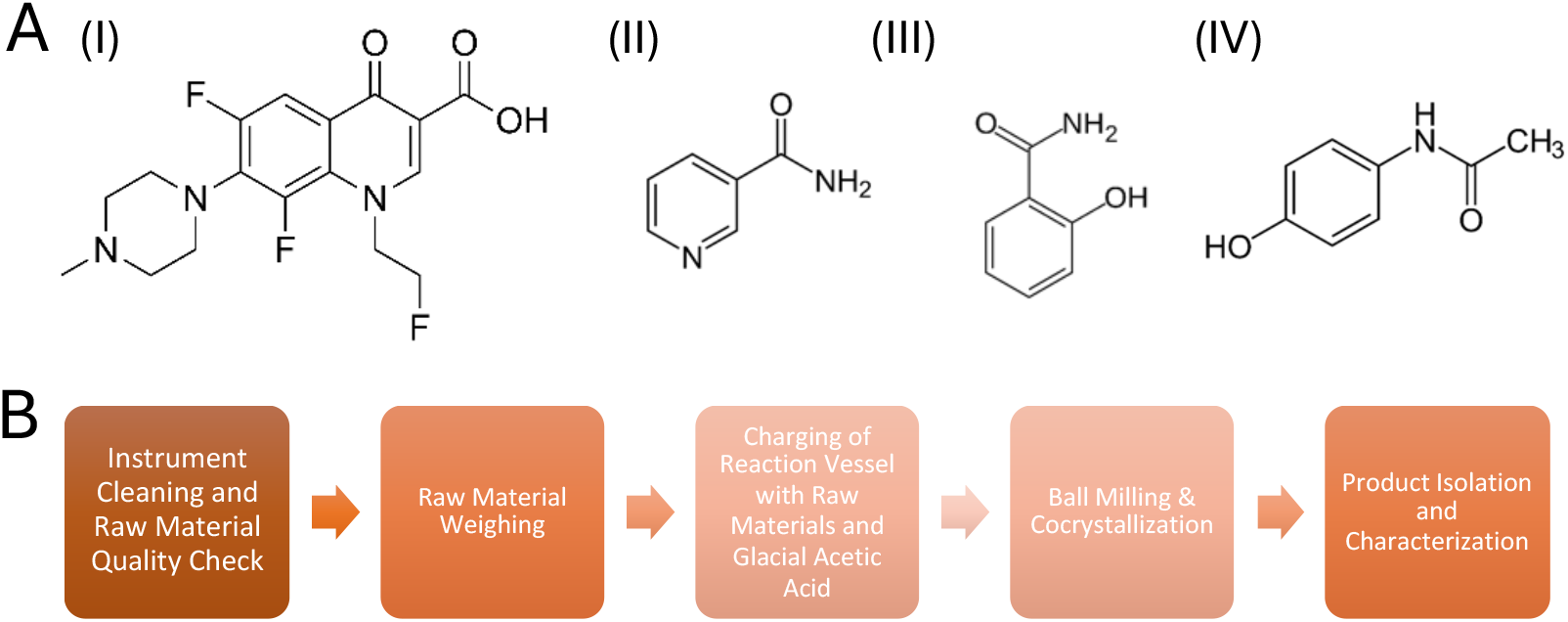
[A] The chemical structures of [I] Fleroxacin, [II] Nicotinamide, [III] Salicylamide, and [IV] Acetaminophen [B] Synthesis scheme for co-crystals.

Our decision regarding the use of glacial acetic acid is partially inspired by the solubility characteristics of Fleroxacin and that there exists a patent utilizing glacial acetic acid as a solvent for pharmaceutical production^19^: Zoladex® an implant containing hormone-retarding peptides which uses glacial acetic acid as a solvent in one of its processes. In its case, the active ingredient (Goserelin) is dissolved in glacial acetic acid prior to compounding with a biodegradable polymer^19–21^. We also have ensured the compatibility of glacial acetic acid by performing dissolution experiments of each raw material separately and recrystallizing them; ensuring no formation of salts as documented by appendix 7.1 and 7.4. In this study, we explore the cocrystallization of Fleroxacin with small molecule APIs and nutrients, namely nicotinamide, salicylamide, and acetaminophen. The selection of these coformers was driven by a combination of trial-and-error screening and the adaptation of a previous report on the cocrystal between Levofloxacin and Metacetamol, published by Taeko Shinozaki from Daiichi Sankyo and Kunikazu Moribe from Chiba University^22^.

We employ a rule called “ΔpK_a_”rule. ΔpK_a_ rule states that “the difference between pK_a_ (ΔpK_a_) of conjugate acid of base – pK_a_ of acid will determine type of products”. Many values reported are actually empirical values based on compilations of results. Therefore, we have chosen rules set by Bhogala *et. al*.^23^ : ΔpK_a_ < 0: no proton transfer between coformer and drug, ΔpK_a_ = [0, 3.75]: cocrystal with salt-cocrystal continuum (treated as cocrystals for our purposes), ΔpK_a_ > 3.75: salt. Experimentally, however, one can also check for proton transfer via a method called X-ray photoelectron spectroscopy (XPS)^24^. A proton that has been from its parent acid to a base will be more difficult to remove and show a higher binding energy than a proton being “shared” between an acid and a base.

Nicotinamide (see figure 1A(II)), a nutrient belonging to the vitamin B complex, offers potential nutritional benefits to patients^25,26^. We hypothesize that we can not only improve the solubility of Fleroxacin but also introduce a nutrient component that may contribute to patient well-being during treatment by cocrystallizing Fleroxacin with nicotinamide. Nicotinamide itself has been used frequently; example is with carbamezapine cocrystal^27^. The pKa value of nicotinamide is approximately 3.35, while that of Fleroxacin is variably reported to be around 5.5 (acidic). At this pH level, the formation of a salt between these compounds is considered unlikely. Additionally, nicotinamide is inherently safe as one must consume a huge amount around 3 g per day for significant toxic effects^26,28^.

Salicylamide (see figure 1A(III)), a drug known for its pain-relieving properties,^29^ has high-water solubility,^30^ and salicylamide serves as a suitable candidate to enhance the solubility of Fleroxacin as the water solubility of salicylamide^31^ also ensures that the resulting cocrystals can readily dissolve, improving the bioavailability and efficacy of Fleroxacin. Furthermore, the pKa value of salicylamide is approximately 8.2. At this pH range, there is a high likelihood of cocrystal formation between these compounds.

Lastly, our choice to investigate the cocrystallization of Fleroxacin with acetaminophen (see figure 1A(IV)) is based on empirical hypothesis and structural similarity, in contrast with the first two. There is a previous report on the cocrystal between Levofloxacin and Metacetamol^22^ (Metacetamol itself is actually an analgesic too and a structural isomer of paracetamol but has never been marketed),^32^ we propose and theorize that a similar cocrystallization principle can be applied to Fleroxacin and paracetamol due to molecular similarities between these two pairs.

## 2. Materials and Methods

All materials were commercially available. Fleroxacin was purchased from AA Blocks Inc. (San Diego, CA, USA). Nicotinamide, salicylamide, paracetamol and glacial acetic acid was purchased from Thermo Fisher Scientific (Waltham, MA, USA). All materials were anhydrous, and all impurities are absent per diffraction data compared to references from Cambridge Structural Database.

### 2.1 Preparation and Synthesis

The sample preparation process was carried out using a Spex® Sample Prep 8000M Mixer mill equipped with a 60 ml steel vessel. To ensure optimal conditions for sample preparation, the vessel underwent a thorough cleaning procedure before each preparation. Firstly, the vessel was handwashed using water and soap to remove any visible contaminants. It was then carefully wiped dry with paper towels to eliminate moisture. Subsequently, the vessel was cleaned with isopropanol and dried again with paper towels to ensure the absence of any residual substances.

For the synthesis of the Fleroxacin cocrystals, an ∼60 ml metal can (sample container) was utilized. Fleroxacin and the corresponding coformer, in a 1:1 molar ratio, were added to the metal vessel. To promote cocrystal formation, several drops of glacial acetic acid were incorporated into each mixture. The use of glacial acetic acid is crucial as cocrystallization without it for our purposes would yield amorphous mixture of starting materials instead of cocrystals.

The specific compositions for the cocrystals are as follows (API & coformer follow 1:1 molar ratio):

#### 2.1.1 Fleroxacin Nicotinamide

The mixture comprised 400 mg of Fleroxacin (1.083 mmol) and 132 mg of Nicotinamide (1.080 mmol), with the addition of 10 drops of glacial acetic acid. During the ball milling process, six metal spheres with a diameter of 1.5 cm were included.

#### 2.1.2 Fleroxacin Salicylamide

This cocrystal formulation consisted of 400 mg of Fleroxacin (1.083 mmol) and 150 mg of Salicylamide (1.093 mmol), combined with 10 drops of glacial acetic acid. Similar to the previous sample, six metal spheres with a diameter of 1.5 cm were employed during ball milling.

#### 2.1.3 Fleroxacin Acetaminophen

For this cocrystal synthesis, 800 mg of Fleroxacin (2.1660) and 327.4 mg of Acetaminophen (2.165 mmol) were mixed with 10 drops of glacial acetic acid. The ball milling process involved the use of four metal spheres with a diameter of 1.5 cm.

The ball milling procedure was conducted for a duration of 30 minutes to ensure thorough mixing and interaction between the components. After completion of the milling process, the resulting material was promptly transferred into 20 ml scintillation vials for storage and subsequent analysis. To prevent the formation of hydrates during the sample preparation, the use of glacial acetic acid was necessary. The drops of glacial acetic acid were accurately dispensed using a 3 ml transfer pipette, ensuring precise control over the volume of acid added to each sample (Figure 1B).

### 2.2 Characterization Techniques

Physically, the samples displayed significant color change, white powder transforming into a yellow-green wet cake that rapidly dried in room temperature and pressure air. X-ray diffraction (XRD) was chosen as the preferred characterization technique because changes in the crystal structure (formation of cocrystal) would directly impact the powder diffraction pattern and by analyzing the XRD patterns, direct information regarding the structural modifications (e.g., due to formation of a new crystal) could be obtained^33^.

The characterization of the synthesized cocrystals mainly relied on X-ray diffraction, for the analysis of all three cocrystals synthesized. Even though per United States Pharmacopeia (USP) <776> Optical Microscopy can indeed be used to identify crystallinity qualities of a sample^34^, (via detecting birefringence/interference colors and extinction positions when the microscope stage is revolved), we use X-ray powder diffraction per USP <941> X-Ray Powder Diffraction due to the ability to see not only if a sample if crystalline but what it is exactly (differentiating mixture vs new powder diffraction pattern).

Elemental analysis (energy dispersive X-ray analysis and Inductively Coupled Plasma Mass Spectrometry/ICP-MS) was considered too for our samples. However, the samples are all organic materials (composed of carbon, hydrogen, oxygen and nitrogen) and our drugs are not heavy metal-containing ones (e.g., cisplatin or Technetium (^99m^Tc) exametazime). Also, other elements other than those mentioned above were not used in the synthesis. Therefore, we have decided to skip this characterization as it does not provide additional insight sufficient enough to justify the procedure). Additionally, per USP <232> Elemental Impurities – Limits and <233> Elemental Impurities – Procedures apply to commercial drug products instead of academic research materials, which are beyond our scope.

Infrared (IR) analysis was excluded as the sizes of individual crystallites can matter a lot due to that this method is dominated by anisotropy. At this stage of our research, particle size controls are beyond our scope and therefore, IR is inapplicable until samples with tightly controlled particle size distributions are made.^35^ Other consideration is that IR analysis deals with absorption in the infrared region, which is dominated by molecular vibrations. Information and metrics this would give include bending, stretching and vibrations of bonds, which are not the most important for crystalline materials and their structures. The last concern in that in USP <197> Spectrophotometric Identification Tests requires the use of a reference standard, which are unknown for these novel cocrystals (commercially unavailable).

Thermal analysis (melting, differential scanning calorimetry are thermogravimetric) were considered but impurity analysis was done via phase purity analysis from the synchrotron data in section 2.6.2.

### 2.3 Powder X-ray Diffraction

We performed standard laboratory X-ray diffraction. To prepare the samples for X-ray diffraction analysis, the materials obtained from the ball milling synthesis were finely ground to break up clumps and prevent any preferred orientation. The sample was then carefully mounted on a nylon loop sample holder using a small amount of mineral oil. X-ray diffraction analysis was conducted using the Rigaku R-Axis Spider instrument with a Curved Image Plate Detector. The experimental geometry was set to transmission mode, and the anode material used was copper (λ=0.15405nm). The X-ray diffraction measurements were performed under the following conditions: a tension of 40 kilovolts, a current of 40 milliamperes, and a collimation of 0.5 mm. The Omega setting was disabled due to the detector type, while the Phi rotation was set at 5 degrees per second. The 2theta range for data collection was set from 2 to 40 degrees.

### 2.4 Synchrotron X-ray Diffraction

To obtain more comprehensive data, rule out operator and instrumentation-derived artifacts, and take advantage of advanced instrumentation, the X-ray diffraction experiment was replicated at Argonne National Laboratory’s Advanced Photon Source in Lemont, IL 60439. The specific beamline used was 11-BM, offering enhanced capabilities. This subsequent analysis at Argonne National Laboratory was motivated by the benefits of synchrotron radiation, which offers virtually perfect monochromatic radiation with exceptionally high brightness. Unlike copper sources, which may be contaminated with k alpha 2 and k beta radiations, synchrotron radiation provides superior spectral purity. Additionally, the advanced optics employed at synchrotron facilities, in comparison to university laboratories, further enhance the accuracy and quality of the XRD data. Lastly, we wished to show that there were no operator-induced errors in that sample preparation and data collection by unrelated and different personnel yield the identical data.

The experimental settings at this facility included a Bending Magnet (BM) as the source whose wavelength was 0.04597 nm. The flux at 30 keV was approximately 5×10^11^ photons per second. The monochromator employed was a Si(111) double crystal in a bounce-up geometry, and the focusing was achieved using a sagittally bent Si(111) crystal and a 1-meter Si/Pt mirror. The beam size at the sample position was 1.5 mm horizontally and 0.5 mm vertically, ensuring a focused beam. The detection system consisted of 12 independent analyzer sets with 2θ separation of approximately 2° using Si(111) analyzer crystals and LaCl_3_ scintillation detectors. The angular coverage spanned a 2θ range of 0.5°-130° at room temperature and pressure temperature, with a maximum Q value (Q_max_) of approximately 28 Å^-1^ at 30 keV. The resolution achieved was ΔQ/Q ≈ 1.4×10^-4^ (minimum 2θ step size = 0.0001°).

The measurements were performed at a temperature of 100 kelvins aiming to minimize thermal disturbances and stabilize the crystal lattice during data collection^36^. Sample preparation was done per the instructions on Argonne National Laboratory’s website, and the actual data collection was done by Argonne staff at the laboratory due to the mail-in service nature of the facility.

### 2.5 Air-sensitive Laboratory X-ray Diffraction (Fleroxacin-Paracetamol Only)

Samples of the Fleroxacin-acetaminophen cocrystal were analyzed at the Texas Material Institute in collaboration with a partner institution, as a synchrotron facility was not accessible for this study. In-house powder diffraction measurements were performed using a specific procedure. The malleable material was spread onto a zero-background Si plate, creating a thin smear of the material. While this method may have introduced some skewness in the recorded intensities, it allowed for a pattern with enhanced angular resolution. To prevent the rapid loss of acetic acid from the specimen, an “air-tight” specimen holder was utilized.

The specimen holder, containing the prepared sample, was then inserted into a Rigaku MiniFlex 600 diffractometer equipped with a Cu anode X-ray (λ=0.15405 nm) tube operating at 40kV/15mA. The diffractometer was fitted with a graphite diffracted beam monochromator to eliminate wavelengths other than Cu-Kα. Patterns were recorded over the 3°-30° 2-theta range, employing a step increment of 0.025° in continuous scan mode with a scanning speed of 0.7°/min. Jade powder pattern processing software from MDI Corporation was used to analyze the data. This software was utilized to determine the positions of peaks in the diffraction patterns. For reference, patterns of standard materials were obtained from the PDF2 database by ICDD Corporation whenever available. In the case of Fleroxacin, patterns were acquired from locally obtained stock starting material, aiming for accurate comparisons and identification of peaks.

### 2.6 Data Analysis

#### 2.6.1 Polymorph Phase Identification

The sample data analysis involved two sets of X-ray diffraction data: one obtained from the low-resolution Rigaku R-Axis Spider system and the other from the high-resolution analysis at Argonne National Lab’s Advanced Photon Source. The X-ray diffraction analysis criteria were based on the guidelines outlined in the United States Pharmacopeia 941 for Characterization of Crystalline and Partially Crystalline Solids by X-Ray Powder Diffraction (XRPD). Specifically, the analysis was conducted within the 2 degrees to 40 degrees two theta angle range, and a peak with a difference of 0.2 two-theta degrees compared to the reference pattern was considered unique (USP <941> X-Ray Powder Diffraction requirement).

The first analysis focused on polymorph phase identification using powder X-ray diffraction with the Rigaku R-Axis Spider system. The resulting powder patterns were compared to the X-ray diffraction patterns of the starting materials: Fleroxacin and its appropriate co-formers. It is important to note that acetic acid, being a liquid, does not exhibit a diffraction pattern, and none of the starting materials can form acetate salts at all. The analysis employed the Bruker EVA software, aiming to identify the emergence of new and distinct diffractogram patterns and the suppression of starting material patterns. The presence of these patterns would indicate the formation of a new cocrystal.

#### 2.6.2 Synchrotron X-ray Confirmation and Phase Purity Analysis

In the subsequent analysis using the synchrotron data obtained from Argonne National Lab’s Advanced Photon Source, the JADE software was employed. The high-resolution data from the synchrotron analysis confirmed the polymorph phase identification results obtained from the Rigaku R-Axis Spider system, thereby validating the previous findings. Furthermore, a “phase identification” analysis was conducted to ensure that the resulting product is a pure cocrystal and not a mixture of unreacted or partially reacted materials (another requirement of cocrystal is that a material must be single-phase)^3^. It should be noted that a minimal amount of unreacted materials may be tolerated, as 100% yield is impossible to achieve, and there is no motivation for that since the FDA tolerates impurities at certain levels anyway^37^.

Worth mentioning is that the United States Pharmacopeia guidelines were adhered to throughout the X-ray diffraction analysis process, ensuring compliance with the established standards for characterizing crystalline and partially crystalline solids. The distinction between “polymorph phase identification” and “phase analysis” lies in their respective objectives. Polymorph phase identification aims to identify different crystal structures (polymorphs) of a compound^38^, while phase analysis focuses on verifying the purity and “single phase-ness” of the material. In this study, both aspects were considered: polymorph phase identification confirmed the formation of a new cocrystal, while phase analysis ensured the purity of the resulting product.

## 3. Results

In Figure 2, the diffractogram clearly demonstrates the presence of a novel material exhibiting a distinct powder diffraction pattern. The marked features, indicated by red arrows, are readily identifiable and provide strong evidence of the material’s unique characteristics. Notably, the peaks corresponding to the starting materials are predominantly suppressed, indicating a significant alteration and integration of these components within the synthesized material. This observation suggests that the starting materials have undergone a transformation, contributing to the formation of the new powder diffraction pattern, it becomes evident that the material’s powder diffraction pattern differs significantly from the starting materials, as the peaks corresponding to these components are diminished in intensity. This suppression of peaks suggests that the starting materials have experienced chemical and structural modifications during the synthesis process. The integration of the starting materials into the new crystal lattice leads to the formation of distinct peaks, indicative of the unique arrangement of atoms within the synthesized material.

**Figure 2:**
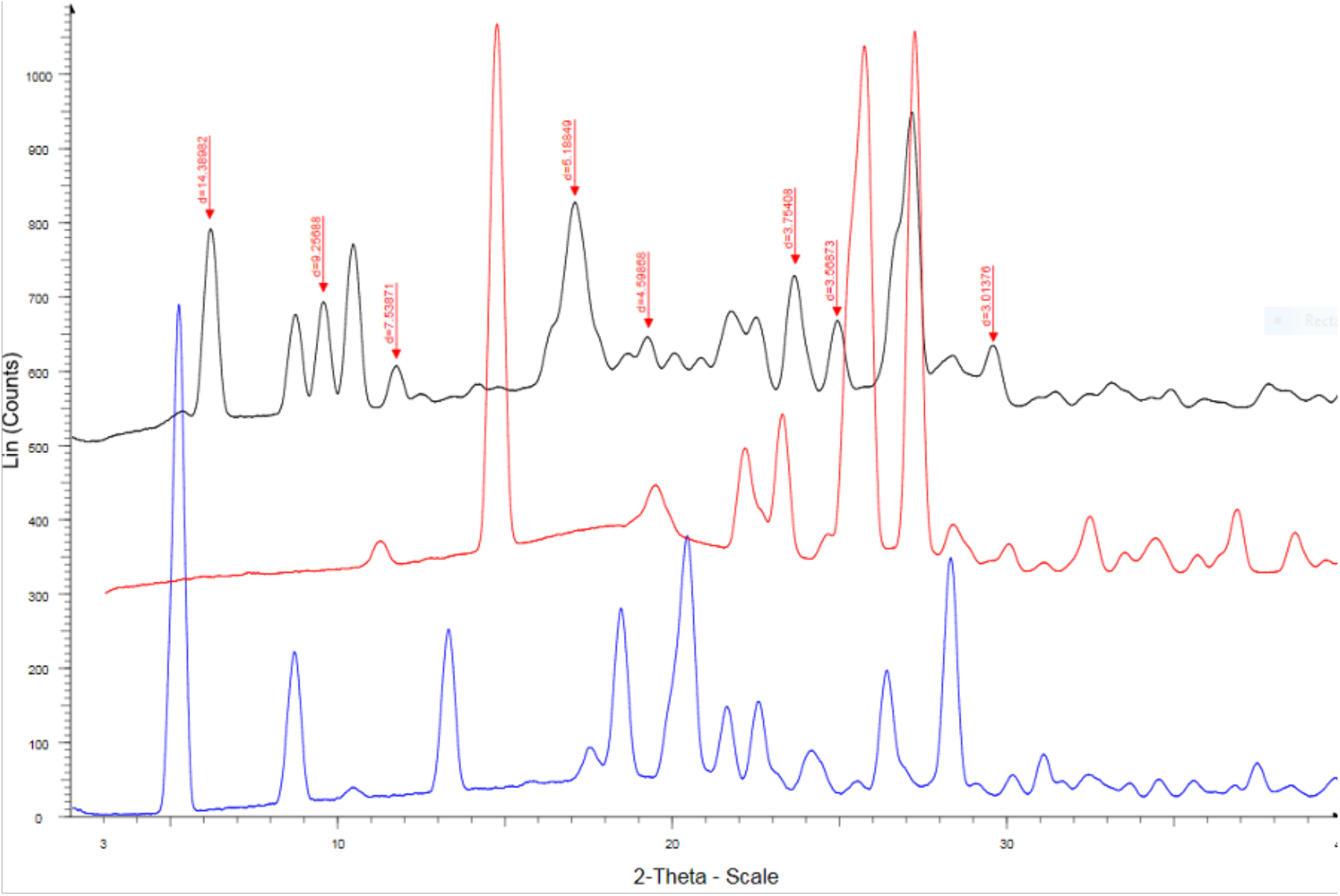
Fleroxacin-Nicotinamide cocrystal from Rigaku R-Axis Spider with copper radiation. Black: product, red: nicotinamide and blue: Fleroxacin. The key peak ‘d’ values were mentioned in red text.

The presence of suppressed peaks in the diffractogram not only confirms the successful synthesis of the new material but also indicates the effective integration of the starting materials into it’s a new cocrystal. This suppression of peaks demonstrates that the synthesized material is not a simple mixture of the starting materials but rather a distinct compound with altered properties. The observed transformations and integration of the starting materials contribute to the formation of a novel cocrystal, giving rise to the unique diffraction pattern observed in the diffractogram. Figure 3 presents the synchrotron diffractogram, which provides compelling evidence of the existence of a new material characterized by a distinct powder diffraction pattern. The compressed two-theta value range (0 to 9°) in this synchrotron experiment, as compared to the Rigaku R-Axis Spider (0 to 40°), is due to the utilization of much shorter wavelengths. However, the fundamental results remain consistent, and if necessary, wavelength conversions can be performed. Notably, clear and easily identifiable peaks in the diffractogram are highlighted with red arrows, indicating the presence of new diffraction patterns from a new material with different crystal structures.

**Figure 3:**
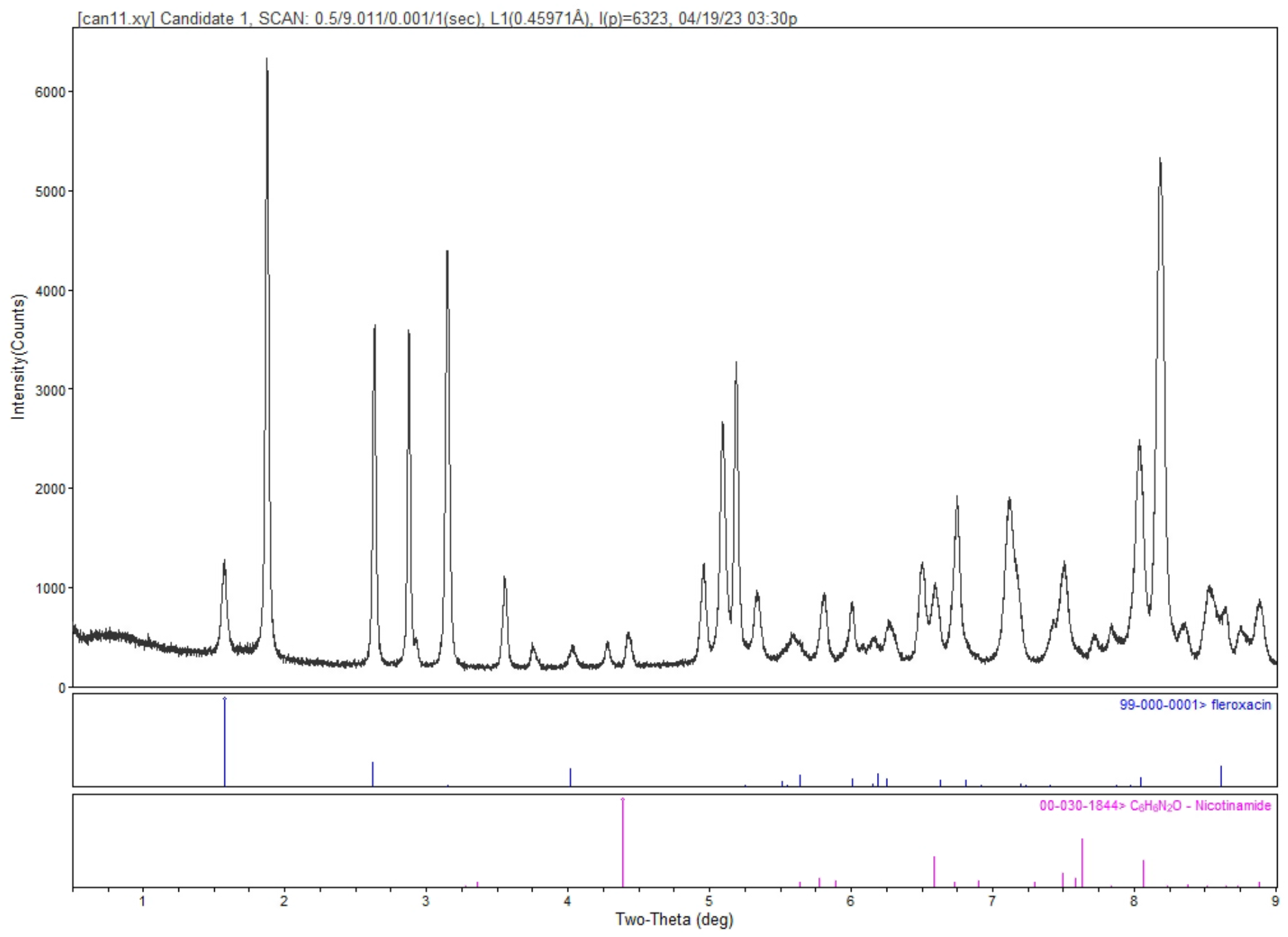
Fleroxacin-Nicotinamide cocrystal from Advanced Photon Source 11-BM synchrotron. Two-theta value is compressed due to the much shorter wavelength used.

Agreement between synchrotron and laboratory X-ray data shows that the peaks are real/not artifacts (e.g., spectral impurity) or instrumental errors (e.g., bad optics or alignment) from the laboratory X-ray diffraction instrument. Secondly, the synchrotron data rules out operator errors-induced artifacts in the laboratory X-ray diffraction data. Lastly, the resulting synchrotron data enables phase analysis (section 2.6.2).

The significant suppression of peaks originating from the starting materials strongly suggests that they have undergone transformations and integrated with some residual components. The distinctiveness of peaks is determined by a 2-theta difference exceeding ±0.2. Importantly, the observed powder diffraction pattern does not indicate the presence of a mere mixture of starting materials, reinforcing the notion that a new material with a unique crystal structure has been formed. Comparing the synchrotron data to the previous results, it becomes evident that the presence of amorphous content, which might be expected from the ball milling process, is absent in the current analysis. This discrepancy can be attributed to the employment of superior post-diffraction optics at the Advanced Photon Source, effectively eliminating artifacts and providing a clearer representation of the powder diffraction pattern. The confirmation of the new material’s existence through the synchrotron data further strengthens the validity of the results obtained. The diffraction patterns obtained from this analysis serve as definitive evidence for the formation of a new cocrystal, underscoring the importance of X-ray diffraction techniques in elucidating the structural characteristics of materials.

Figure 4 exhibits a diffractogram that unequivocally demonstrates the presence of a novel material characterized by a distinct powder diffraction pattern. Prominent features, conveniently identified with manually placed blue arrows, highlight the characteristic peaks indicative of the crystal lattice arrangement. The suppression of peaks associated with the starting materials suggests their transformation and incorporation into the cocrystal structure. Notably, pairs of peaks with a 2-theta difference exceeding ±0.2° are deemed distinguishable, emphasizing the distinct nature of the materia^38^. Importantly, the diffractogram confirms that it is not a mere mixture of the starting materials. Similar to the previous cocrystal pair, where the presence of apparent amorphous content was revealed to be an artifact caused by the absence of post-diffraction optics, our expectations align with a comparable scenario for the material under investigation. The initial analysis, indicating significant amorphous content, is likely a result of limitations in the experimental setup and the absence of advanced post-diffraction optics. To ascertain the true nature of the material, further examination employing synchrotron-based X-ray diffraction was conducted.

**Figure 4:**
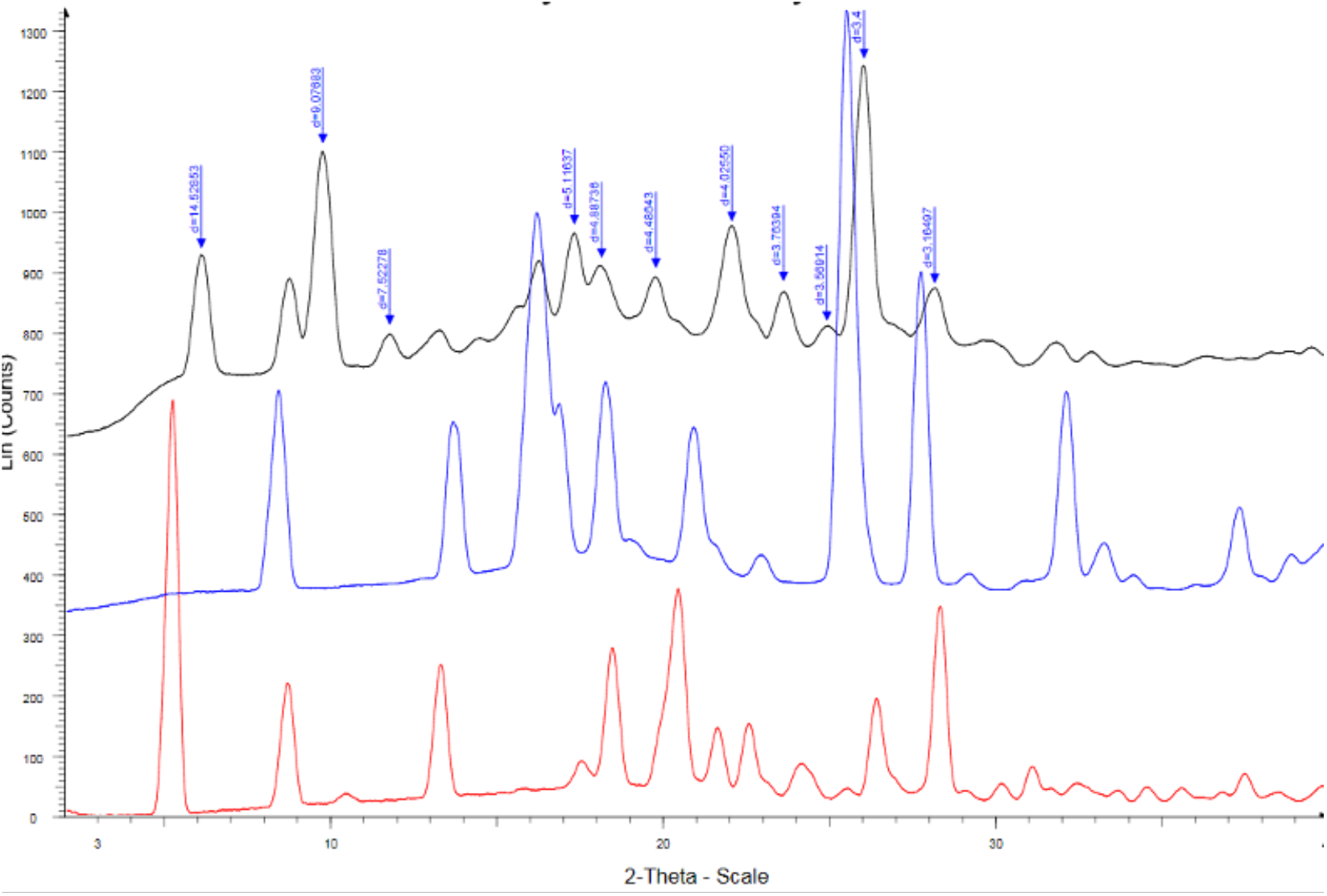
Fleroxacin-Salicylamide cocrystal from Rigaku R-Axis Spider with copper radiation. Black: product, blue: salicylamide, and red: Fleroxacin.

The analysis of this data set follows a similar pattern to the previous cocrystal examination. The synchrotron diffractogram presented in Figure 5 unambiguously demonstrates the presence of a novel material with a distinct powder diffraction pattern, thereby supporting the earlier findings obtained using the Rigaku R-Axis Spider. The two-theta value range of this synchrotron analysis (0 to 9°) is compressed due to the utilization of a much shorter wavelength, but the fundamental results remain consistent, and wavelength conversion is possible if needed. Prominent features are evidently identified and highlighted with manual red arrows. The suppression of peaks corresponding to the starting materials indicates their transformation and integration, with some remnants remaining. In accordance with our criteria, pairs of peaks are considered distinct if the difference in their two-theta values exceeds ±0.2°. Importantly, the diffractogram is not indicative of a mere mixture of the initial materials. Comparing the synchrotron data with the previous results, it becomes evident that the apparent presence of amorphous content, which was observed in the earlier analysis, is now absent. This discrepancy can be attributed to the implementation of superior post-diffraction optics at the Advanced Photon Source, effectively eliminating the artifacts that previously misled our interpretation. The absence of amorphous signals in the synchrotron data further strengthens the validity of the findings.

**Figure 5:**
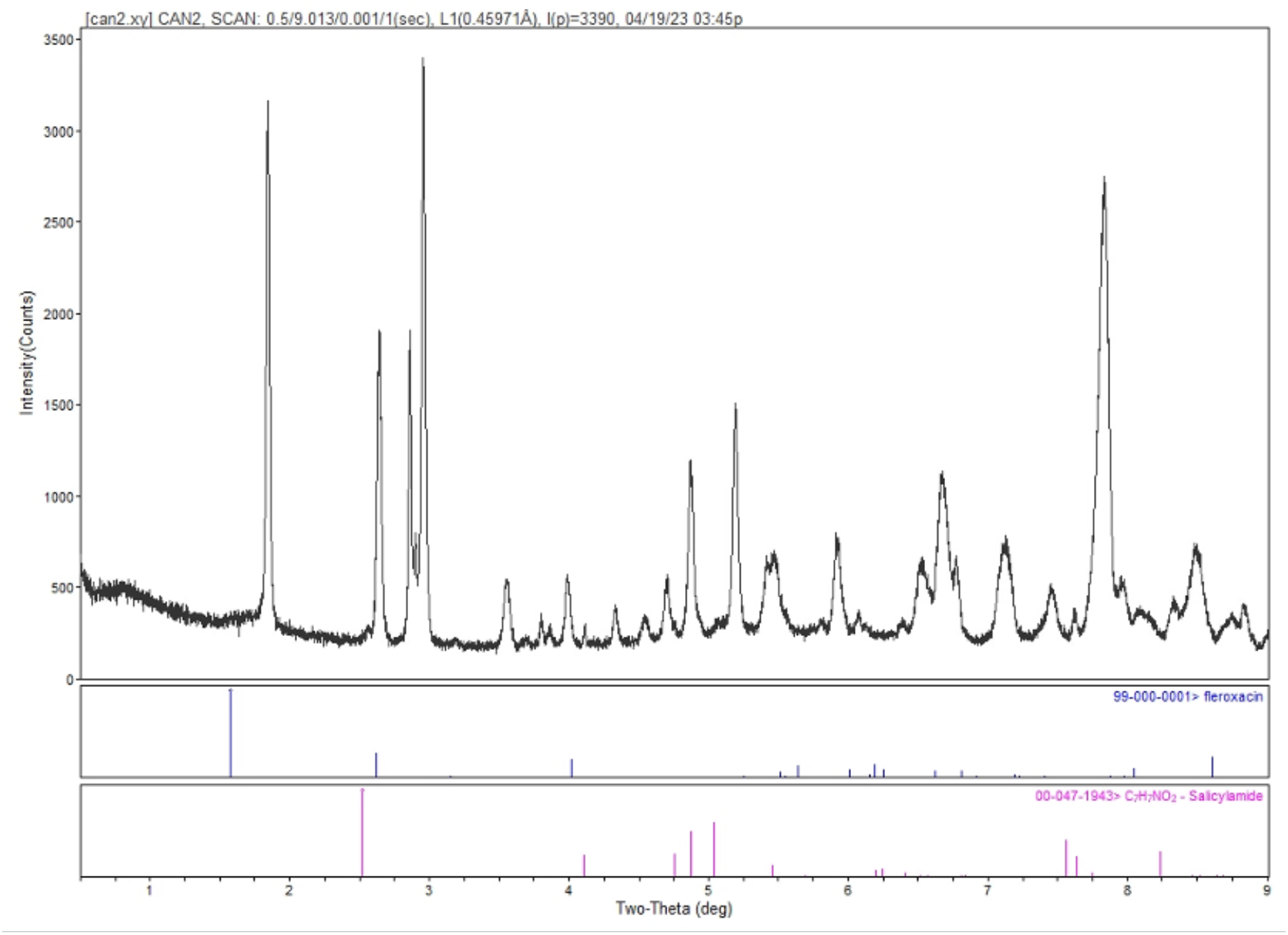
Fleroxacin-Salicylamide cocrystal from Advanced Photon Source 11-BM synchrotron. Two-theta value is compressed due to the much shorter wavelength used.

The synchrotron data is identical and shows agreement to the laboratory X-ray diffraction data, showing that the peaks are real (identical compared to the laboratory X-ray diffraction instrument’s). Conclusively, the synchrotron data obtained confirms the existence of a new cocrystal as indicated by the diffraction patterns (and rules out instrument artifacts or preparation errors). The distinct diffraction patterns of the material is obviously established, reinforcing the robustness and accuracy of our results.

Figure 6 displays the diffractogram obtained using a Miniflex instrument (as a synchrotron was not available), revealing the presence of a distinct powder diffraction pattern for the new material. Notably identifiable peaks are highlighted with red arrows, indicating the formation of a new compound. The majority of peaks from the starting materials are suppressed, indicating their transformation and integration with small amounts of unreacted components. Distinct peaks are characterized by a 2-theta difference beyond ±0.2°, further confirming that the diffractogram does not originate from a mere mixture of starting materials. In the obtained pattern, there are no discernible Fleroxacin peaks; however, acetaminophen peaks are detectable along with a series of other peaks. This observation suggests a stoichiometric effect, indicating that there are likely two Fleroxacin molecules for each acetaminophen molecule in the cocrystal. Despite the use of a non-synchrotron source, the inclusion of post-diffraction optics has aided in minimizing artifacts. The data obtained demonstrates that the product exhibits a high degree of crystallinity with minimal presence of amorphous materials.

**Figure 6:**
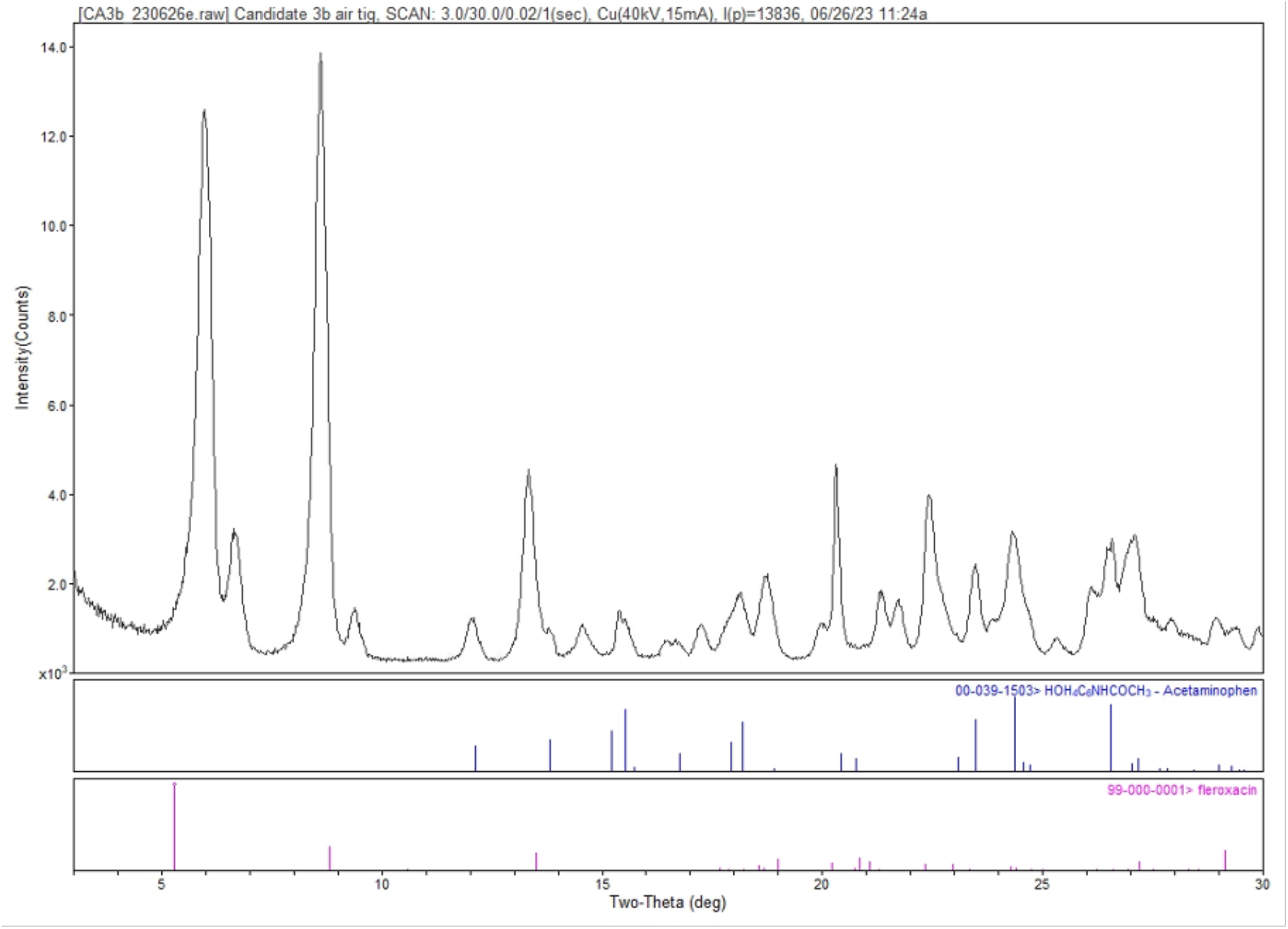
Fleroxacin-Acetaminophen cocrystal from Rigaku MiniFlex. This one is collected with X-rays from a copper anode (benchtop lab-scale).

These findings highlight the successful formation of a new crystalline material through the cocrystallization process. The integration of Fleroxacin and acetaminophen has resulted in the formation of a distinct compound with a well-defined powder diffraction pattern. The use of post-diffraction optics has further ensured the accuracy and reliability of the obtained data, providing valuable insights into the stoichiometry and crystallinity of the cocrystal.

## 4. Discussion

The major finding of this study is the creation of novel and unexpected Fleroxacin cocrystals with three nutrients and over-the-counter small molecule drugs/active pharmaceutical ingredients (APIs). This fact alone is very interesting given the difficulties of cocrystallizing Fleroxacin despite its favorable chemical structure and presence of various hydrogen bonding-supportive functional groups (e.g., carboxylic acid). These results are also empirical proof of the delta pKa rule used to predict if a pair of materials can form cocrystals. As noted before, the pKas of each component are known, and feeding them into the formulas can describe whether the formation of cocrystals is theoretically possible. Additionally, our results are consistent with other papers in terms of powder X-ray diffractogram behaviors: the creation of a new crystal structure will suppress the peaks of the starting materials while showing new peaks; the total number of peaks will be different.

The only intriguing part is the Fleroxacin-acetaminophen diffractogram, in that residual peaks of one of the starting materials are present. This could mean either that the new material has a non-1:1 (drug:coformer) ratio or something else (e.g., failure to cocrystallize). We believe it’s the former (non-1:1 cocrystal) since neither Fleroxacin nor acetaminophen forms salts with acetic acid, while the new peaks are clearly distinct. The other major discovery is the use of glacial acetic acid as a way to solubilize and catalyze cocrystallization of pharmaceuticals. Despite its high dissolving abilities, glacial acetic acid is seldom used as a catalyst or solvent for cocrystallization. There are reports of its use combined with other reagents, but not purely by itself. Zoladex® is a notable case, but it even formed acetate salts. In our discovery, we are also fortunate that none of our reagents formed stable acetate salts with glacial acetic acid. For any readers would like to know more about this, see figure S1, S2 and S5 in the supporting information, supporting information section 1.1 addresses Fleroxacin & nicotinamide while section 1.4 address salicylamide. Paracetamol (structure: phenol with N-acetyl group) is not known to form acetate salts.

Of course, all of our products were confirmed with both university laboratory-grade X-ray diffraction machines and a synchrotron X-ray source to ensure purity and verify the creation of a new molecular entity (also, identical data from synchrotron and lab X-ray data disprove possibilities of instrumentation, spectral and human error artifacts). The current limitation is that the method used to create the cocrystals is inherently hostile to the growth of large single crystals (and even as long as 6 months)^39^. This rules out the use of traditional single-crystal X-ray diffraction apparatus to probe the actual structure of the unit cell. The easiest way to remedy this, however, is to use electron beam diffraction (commonly known as MicroED), which allows the use of nanometer/micrometer-sized (literally a billionth-sized crystal can be used) samples to perform single-crystal diffraction^40,41^.

## 5. Conclusion

In this study, we successfully formed novel cocrystals of Fleroxacin with nicotinamide, salicylamide, and acetaminophen using a novel acetic acid catalyst. X-ray diffraction (XRD) confirmed the cocrystal formation through distinctive patterns and physical appearance changes. These findings have significant implications for improving Fleroxacin’s properties and therapeutic efficacy. Synchrotron (and university lab) powder XRD played a crucial role in determining the powder diffraction pattern and understanding the molecular arrangement within the cocrystal lattice, guiding future formulation strategies. Phase identification analyses ensured cocrystal purity, and the use of post-diffraction optics accurately assessed amorphous content. Moreover, the successful synthesis and characterization of these Fleroxacin cocrystals with salicylamide, nicotinamide, and acetaminophen with uncommonly used yet perfectly edible catalyst (acetic acid) should further bolster green chemistry in that use of green and safe catalyst can start as early as bench scale and potentially survive as far as industrial scale. The use of green and safe catalyst can also simply quality control in the future. While limitations exist, further exploration of characterization techniques is recommended for a comprehensive understanding. This research advances pharmaceutical development and has the potential to provide enhanced options for patients.

## 6. Acknowledgements

The authors thank the Argonne National Laboratory’s Advanced Photon Source in Lemont, IL 60439 for providing access to use the synchrotron for this study. Authors also acknowledge the Texas Materials Institute at The University of Texas at Austin for central facility to access the state-of-the-art X-ray Diffractometry.

